# The genetic architecture of target-site resistance to pyrethroid insecticides in the African malaria vectors *Anopheles gambiae* and *Anopheles coluzzii*

**DOI:** 10.1101/323980

**Authors:** Chris S. Clarkson, Alistair Miles, Nicholas J. Harding, David Weetman, Dominic Kwiatkowski, Martin Donnelly, The *Anopheles gambiae* 1000 Genomes Consortium

## Abstract

Resistance to pyrethroid insecticides is a major concern for malaria vector control, because these are the compounds used in almost all insecticide-treated bed-nets (ITNs), and are also widely used for indoor residual spraying (IRS). Pyrethroids target the voltage-gated sodium channel (VGSC), an essential component of the mosquito nervous system, but substitutions in the amino acid sequence can disrupt the activity of these insecticides, inducing a resistance phenotype. Here we use Illumina whole-genome sequence data from phase 1 of the *Anopheles gambiae* 1000 Genomes Project (Ag1000G) to provide a comprehensive account of genetic variation in the *Vgsc* gene in mosquito populations from eight African countries. In addition to the three known resistance alleles, we describe 20 non-synonymous nucleotide substitutions at appreciable frequency in one or more populations that are previously unknown in *Anopheles* mosquitoes. Thirteen of these novel alleles were found to occur almost exclusively on haplotypes carrying the known `L995F` resistance allele (L1014F in *Musca domesticus* codon numbering), and may enhance or compensate for the `L995F` resistance pheno-type. A novel mutation I1527T, which is adjacent to a predicted pyrethroid binding site, was found in tight linkage with either of two alleles causing a V402L substitution, similar to a combination of substitutions found to cause pyrethroid resistance in several other insect species. We analyse the genetic backgrounds on which non-synonymous alleles are found, to determine which alleles have experienced recent positive selection, and to refine our understanding of the spread of resistance between species and geographical locations. We describe twelve distinct haplotype groups with evidence of recent positive selection, five of which carry the known `L995F` resistance allele, five of which carry the known `L995S` resistance allele, one of which carries the novel I1527T allele, and one of which carries a novel M490I allele. Seven of these groups are localised to a single geographical location, and five comprise haplotypes from different countries, in one case separated by over 3000 km, providing new information about the geographical distribution and spread of resistance. We also find evidence for multiple introgression events transmitting resistance alleles between *An. gambiae* and *An. coluzzii*. We identify markers that could be used to design high-throughput, low-cost genetic assays for improved surveillance of pyrethroid resistance in the field. Our results demonstrate that the molecular basis of target-site pyrethroid resistance in malaria vectors is more complex than previously appreciated, and provide a foundation for the development of new genetic tools to track the spread insecticide resistance and improve the design of strategies for insecticide resistance management.

## Introduction

Pyrethroid insecticides have been the cornerstone of malaria prevention in Africa for almost two decades [1]. Pyrethroids are currently used in all insecticide-treated bed-nets (ITNs), and are widely used in indoor residual spraying (IRS) campaigns as well as in agriculture. Pyrethroid resistance is widespread in malaria vector populations across Africa [2]. The World Health Organization (WHO) has published plans for insecticide resistance management (IRM), which emphasise the need for improvements in both our knowledge of the molecular mechanisms of resistance and our ability to monitor them in natural populations [3].

The voltage-gated sodium channel (VGSC) is the physiological target of pyrethroid insecticides, and is integral to the insect nervous system. Pyrethroid molecules bind to sites within the protein channel and prevent normal nervous system function, causing paralysis (“knock-down”) and then death. However, amino acid substitutions at key positions within the protein alter the interaction with insecticide molecules (target-site resistance), increasing the dose of insecticide required for knock-down [4, 5]. In the African malaria vectors *Anopheles gambiae* and *An. coluzzii*, three substitutions have been found to cause pyrethroid resistance. Two of these substitutions occur in codon 995^1^, with `L995F` prevalent in West and Central Africa [6, 7], and `L995S` found in Central and East Africa [8, 7]. A third substitution, `N1570Y`, has been found in West and Central Africa and shown to increase resistance in association with `L995F` [10]. However, studies in other insect species have found a variety of other *Vgsc* substitutions inducing a resistance phenotype [11, 12, 5]. To our knowledge, no studies in malaria vectors have analysed the full *Vgsc* coding sequence, thus the molecular basis of target-site resistance to pyrethroids has not been fully explored.

Basic information is also lacking about the spread of pyrethroid resistance in malaria vectors. For example, it is not clear when, where or how many times pyrethroid target-site resistance has emerged. Geographical paths of transmission, carrying resistance alleles between mosquito populations, are also not known. Previous studies have found evidence that `L995F` occurs on several different genetic backgrounds, suggesting multiple independent outbreaks of resistance driven by this allele [13, 14, 15, 16]. However, these studies analysed only small gene regions in a limited number of mosquito populations, and therefore had limited resolution to make inferences about relationships between haplotypes carrying this allele. It has also been shown that the `L995F` allele spread from *An. gambiae* to *An. coluzzii* in West Africa [17, 18]. However, both `L995F` and `L995S` now have wide geographical distributions [7], and to our knowledge no attempts have been made to infer or track the geographical spread of either allele.

Here we report an in-depth analysis of genetic variation in the *Vgsc* gene, using whole-genome Illumina sequence data from phase 1 of the *Anopheles gambiae* 1000 Genomes Project (Ag1000G) [19]. The Ag1000G phase 1 resource includes data on nucleotide variation in 765 wild-caught mosquitoes sampled from 8 countries, with representation of West, Central, Southern and East Africa, and of both *An. gambiae* and *An. coluzzii*. We investigate variation across the complete gene coding sequence, and report population genetic data for both known and novel non-synonymous nucleotide substitutions. We then use haplotype data from the chromosomal region spanning the *Vgsc* gene to study the genetic backgrounds carrying resistance alleles, infer the geographical spread of resistance between mosquito populations, and provide evidence for recent positive selection. Finally, we explore ways in which variation data from Ag1000G can be used to design high-throughput, low-cost genetic assays for surveillance of pyrethroid resistance, with the capability to differentiate and track resistance outbreaks.

## Results

### *Vgsc* non-synonymous nucleotide variation

To identify variants with a potentially functional role in pyrethroid resistance, we extracted single nucleotide polymorphisms (SNPs) that alter the amino acid sequence of the VGSC protein from the Ag1000G phase 1 data resource. We then computed their allele frequencies among 9 mosquito populations defined by species and country of origin. Alleles that confer resistance are expected to increase in frequency under selective pressure, and we filtered the list of potentially functional variant alleles to retain only those at or above 5% frequency in one or more populations (Table 1). The resulting list comprises 23 variant alleles, including the known `L995F`, `L995S` and `N1570Y` resistance alleles, and a further 20 alleles not previously described in anopheline mosquitoes. We reported 15 of these novel alleles in our overall analysis of the Ag1000G phase 1 data resource [19], and we extend the analyses here to incorporate a SNP which alters codon 1603 and two tri-allelic SNPs affecting codons 402 and 490.

The two known resistance alleles affecting codon 995 had the highest overall allele frequencies within the Ag1000G phase 1 cohort (Table 1). The `L995F` allele was at high frequency in populations of both species from West, Central and Southern Africa. The `L995S` allele was at high frequency among *An. gambiae* populations from Central and East Africa. Both alleles were present in *An. gambiae* populations sampled from Cameroon and Gabon, including some individuals with a hybrid `L995F`/S genotype (46/275 individuals in Cameroon, 36/56 in Gabon). In Cameroon these alleles were in Hardy Weinberg equilibrium (*χ*^2^ = 0.02, P > 0.05), but there was an excess of heterozygotes in Gabon (*χ*^2^= 8.96, P < 0.005), suggesting a fitness advantage for mosquitoes carrying both alleles at least in some circumstances.

**Table 1.** Non-synonymous nucleotide variation in the voltage-gated sodium channel gene. AO=Angola; BF=Burkina Faso; GN=Guinea; CM=Cameroon; GA=Gabon; UG=Uganda; KE=Kenya; GW=Guinea-Bissau; *Ac*=*An. coluzzii*; *Ag*=*An. gambiae*. Species status of specimens from Kenya and Guinea-Bissau is uncertain [19]. All variants are at 5% frequency or above in one or more of the 9 Ag1000G phase 1 populations, with the exception of 2,400,071 G>T which is only found in the CM*Ag* population at 0.4% frequency but is included because another mutation (2,400,071 G>A) is found at the same position causing the same amino acid substitution (M490I); and 2,431,019 T>C (F1920S) which is at 4% frequency in GA*Ag* but also found in CM*Ag* and linked to `L995F`.

| Position<sup>1</sup> | Variant |  |  | Population allele frequency (%) |  |  |  |  |  |  |  |  |
| --- | --- | --- | --- | --- | --- | --- | --- | --- | --- | --- | --- | --- |
|  | *Ag*<sup>2</sup> | *Md*<sup>3</sup> | Domain<sup>4</sup> | AOAc | BFAC | GNAg | BFAG | CMAg | GAAg | UGAg | KE | GW |
| 2,390,177 G>A | R254K | R261 | IL45 | 0 | 0 | 0 | 0 | 32 | 21 | 0 | 0 | 0 |
| 2,391,228 G>C | V402L | V410 | IS6 | 0 | 7 | 0 | 0 | 0 | 0 | 0 | 0 | 0 |
| 2,391,228 G>T | V402L | V410 | IS6 | 0 | 7 | 0 | 0 | 0 | 0 | 0 | 0 | 0 |
| 2,399,997 G>C | D466H | - | LI/II | 0 | 0 | 0 | 0 | 7 | 0 | 0 | 0 | 0 |
| 2,400,071 G>A | M490I | M508 | LI/II | 0 | 0 | 0 | 0 | 0 | 0 | 0 | 18 | 0 |
| 2,400,071 G>T | M490I | M508 | LI/II | 0 | 0 | 0 | 0 | 0 | 0 | 0 | 0 | 0 |
| 2,416,980 C>T | T791M | T810 | IIS1 | 0 | 1 | 13 | 14 | 0 | 0 | 0 | 0 | 0 |
| 2,422,651 T>C | L995S | L1014 | IIS6 | 0 | 0 | 0 | 0 | 15 | 64 | 100 | 76 | 0 |
| 2,422,652 A>T | L995F | L1014 | IIS6 | 86 | 85 | 100 | 100 | 53 | 36 | 0 | 0 | 0 |
| 2,424,384 C>T | A1125V | K1133 | LII/III | 9 | 0 | 0 | 0 | 0 | 0 | 0 | 0 | 0 |
| 2,425,077 G>A | V1254I | I1262 | LII/III | 0 | 0 | 0 | 0 | 0 | 0 | 0 | 0 | 5 |
| 2,429,617 T>C | I1527T | I1532 | IIIS6 | 0 | 14 | 0 | 0 | 0 | 0 | 0 | 0 | 0 |
| 2,429,745 A>T* | N1570Y | N1575 | LIII/IV | 0 | 26 | 10 | 22 | 6 | 0 | 0 | 0 | 0 |
| 2,429,897 A>G | E1597G | E1602 | LIII/IV | 0 | 0 | 6 | 4 | 0 | 0 | 0 | 0 | 0 |
| 2,429,915 A>C | K1603T | K1608 | IVS1 | 0 | 5 | 0 | 0 | 0 | 0 | 0 | 0 | 0 |
| 2,430,424 G>T | A1746S | A1751 | IVS5 | 0 | 0 | 11 | 13 | 0 | 0 | 0 | 0 | 0 |
| 2,430,817 G>A | V1853I | V1858 | COOH | 0 | 0 | 8 | 5 | 0 | 0 | 0 | 0 | 0 |
| 2,430,863 T>C | I1868T | I1873 | COOH | 0 | 0 | 18 | 25 | 0 | 0 | 0 | 0 | 0 |
| 2,430,880 C>T | P1874S | P1879 | COOH | 0 | 21 | 0 | 0 | 0 | 0 | 0 | 0 | 0 |
| 2,430,881 C>T | P1874L | P1879 | COOH | 0 | 7 | 45 | 26 | 0 | 0 | 0 | 0 | 0 |
| 2,431,019 T>C | F1920S | Y1925 | COOH | 0 | 0 | 0 | 0 | 1 | 4 | 0 | 0 | 0 |
| 2,431,061 C>T | A1934V | A1939 | COOH | 0 | 12 | 0 | 0 | 0 | 0 | 0 | 0 | 0 |
| 2,431,079 T>C | I1940T | I1945 | COOH | 0 | 4 | 0 | 0 | 7 | 0 | 0 | 0 | 0 |

<sup>1</sup> Position relative to the AgamP3 reference sequence, chromosome arm 2L. Variants marked with an asterisk (\*) failed conservative variant filters applied genome-wide in the Ag1000G phase 1 AR3 callset, but appeared sound on manual inspection of read alignments.
<sup>2</sup> Codon numbering according to *Anopheles gambiae* transcript AGAP004707-RA in geneset AgamP4.4.
<sup>3</sup> Codon numbering according to *Musca domestica* EMBL accession X96668 [9].
<sup>4</sup> Location of the variant within the protein structure. Transmembrane segments are named according to domain number (in Roman numerals) followed by 'S' then the number of the segment; e.g., 'IIS6' means domain two, transmembrane segment six. Internal linkers between segments within the same domain are named according to domain (in Roman numerals) followed by 'L' then the numbers of the linked segments; e.g., 'IL45' means domain one, linker between transmembrane segments four and five. Internal linkers between domains are named 'L' followed by the linked domains; e.g., 'LI/II' means the linker between domains one and two. 'COOH' means the internal carboxyl tail.

The `N1570Y` allele was present in Guinea, Burkina Faso (both species) and Cameroon. This allele has only ever been found in association with `L995F` [10], and has been shown to substantially increase pyrethroid resistance when it occurs in combination with `L995F`, both in association tests of phenotyped field samples [10] and experimentally [20]. To study the patterns of association among non-synonymous variants, we used haplotypes from the Ag1000G phase 1 resource to compute the normalised coefficient of linkage disequilibrium (*D^/^*) between all pairs of variant alleles (Figure 1). As expected, we found `N1570Y` in almost perfect linkage with `L995F`, meaning that `N1570Y` was almost only ever found on haplotypes carrying `L995F`. Of the 20 novel non-synonymous alleles, 13 also occurred almost exclusively in combination with `L995F`, exhibiting the same LD pattern as `N1570Y` (Figure 1). These included two variants in codon 1874 (`P1874S`, `P1874L`), one of which (`P1874S`) has previously been associated with pyrethroid resistance in the crop pest *Plutella xylostella* [21]. The abundance of high-frequency non-synonymous variants occurring in combination with `L995F` is striking for two reasons. First, *Vgsc* is a highly conserved gene, expected to be under strong functional constraint and therefore purifying selection, and so any non-synonymous variants are expected to be rare [11]. Second, in contrast with `L995F`, we did not observe any high-frequency non-synonymous variants occuring in combination with L995S. This contrast was highly significant when data on all variants within the gene were considered: relative to haplotypes carrying the wild-type L995 allele, the ratio of non-synonymous to synonymous nucleotide diversity (*π_N_ /π_S_*) was 28.1 (95% CI [25.2, 31.2]) times higher among haplotypes carrying `L995F` but 1.5 (95% CI [0.8, 2.2]) times higher among haplotypes carrying L995S. These results indicate that `L995F` has substantially altered the selective regime for other amino acid positions within the protein. A number of secondary substitutions have occurred and risen in frequency, pressure.

**Figure 1.**
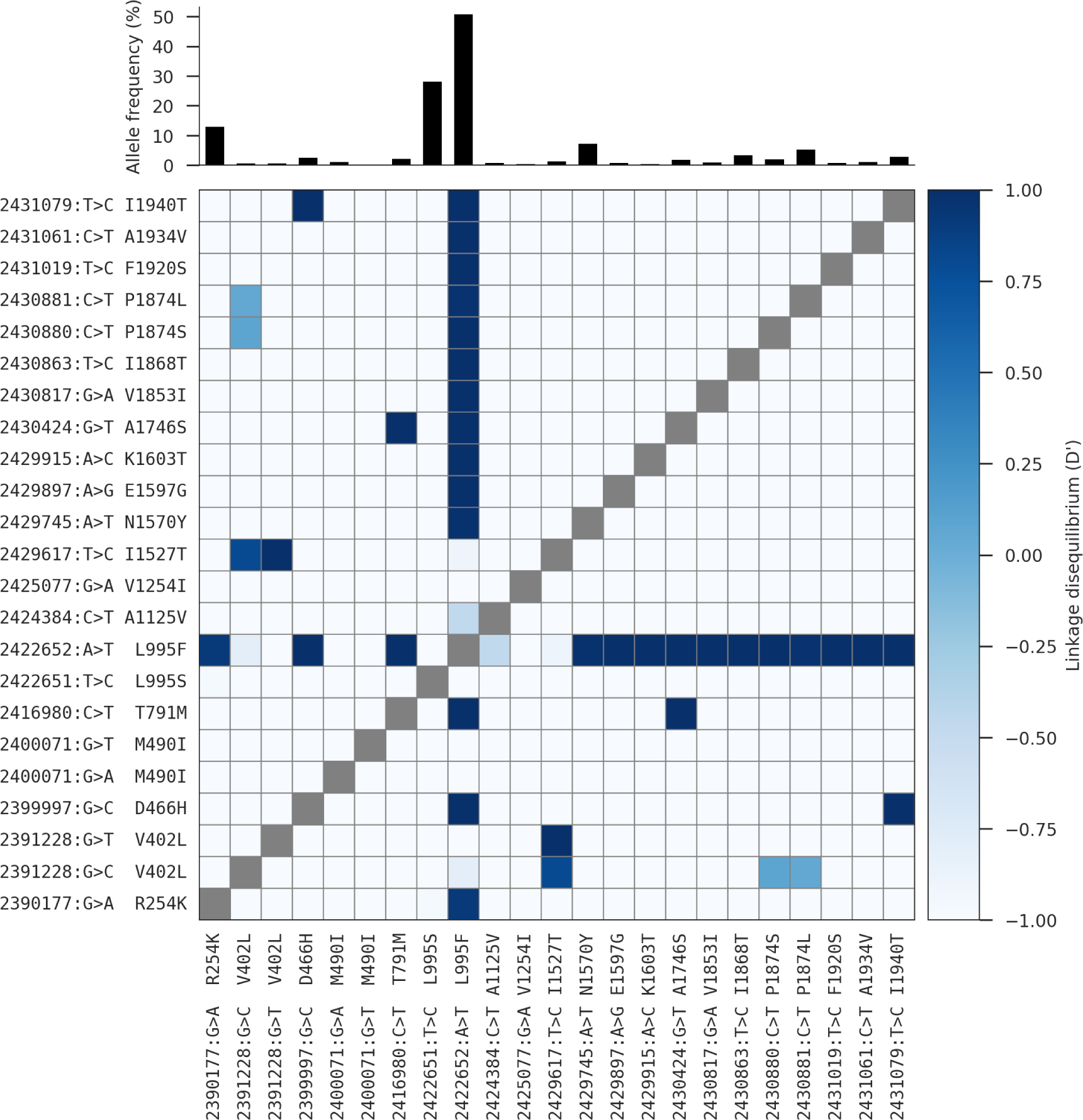
Linkage disequilibrium (*D′*) between non-synonymous variants. A value of 1 indicates that two alleles are in perfect linkage, meaning that one of the alleles is only ever found in combination with the other. Conversely, a value of −1 indicates that two alleles are never found in combination with each other. The bar plot at the top shows the frequency of each allele within the Ag1000G phase 1 cohort. See Table 1 for population allele frequencies.

A novel allele, `I1527T`, was present in *An. coluzzii* from Burkina Faso at 14% frequency. Codon 1527 occurs within trans-membrane segment `IIIS6`, immediately adjacent to residues within a predicted binding site for pyrethroid molecules, thus it is plausible that `I1527T` could alter pyrethroid binding [22, 5]. We also found that the two variant alleles affecting codon 402, both of which induce a `V402L` substitution, were in strong linkage with `I1527T` (*D^/^ ≥* 0.8; Figure 1), and almost all haplotypes carrying `I1527T` also carried a `V402L` substitution. Substitutions in codon 402 have been found in a number of other insect species and shown experimentally to confer pyrethroid resistance [5]. Because of the limited geographical distribution of these alleles, we hypothesize that the `I1527T+V402L` combination represents a pyrethroid resistance allele that arose in West African *An. coluzzii* populations. However, the `L995F` allele is at higher frequency (85%) in our Burkina Faso *An. coluzzii* population, and is known to be increasing in frequency [23], therefore `L995F` may provide a stronger resistance phenotype and is replacing `I1527T+V402L`.

The remaining 4 novel alleles (two separate nucleotide substitutions causing `M490I; A1125V; V1254I`) did not occur in combination with any known resistance allele (Table 1). All are private to a single population, and to our knowledge none have previously been found in other species [12, 5].

### Genetic backgrounds carrying resistance alleles

The Ag1000G data resource provides a rich source of information about the spread of insecticide resistance alleles in any given gene, because data are available not only for SNPs in protein coding regions, but also SNPs in introns and flanking intergenic regions, and in neighbouring genes. These additional variants can be used to analyse the genetic backgrounds (haplotypes) on which resistance alleles are found. In our initial report of the Ag1000G phase 1 resource [19], we used 1710 biallelic SNPs from within the 73.5 kbp *Vgsc* gene (1607 intronic, 103 exonic) to compute the number of SNP differences between all pairs of 1530 haplotypes derived from 765 wild-caught mosquitoes. We then used pairwise genetic distances to perform hierarchical clustering, and found that haplotypes carrying resistance alleles in codon 995 were grouped into 10 distinct clusters, each with near-identical haplotypes. Five of these clusters contained haplotypes carrying the `L995F` allele (labelled F1-F5), and a further five clusters contained haplotypes carrying `L995S` (labelled S1-S5).

To further investigate genetic backgrounds carrying resistance alleles, we used the Ag1000G haplotype data to construct median-joining networks [24] (Figure 2). The network analysis improves on hierarchical clustering by allowing for the reconstruction and placement of intermediate haplotypes that may not be observed in the data. It also allows for non-hierarchical relationships between haplotypes, which may arise if recombination events have occured between haplotypes. We constructed the network up to a maximum edge distance of 2 SNP differences, to ensure that each connected component captures a group of closely-related haplotypes. The resulting network contained 5 groups containing haplotypes carrying `L995F`, and a further 5 groups carrying L995S, in close correspondence with previous results from hierarchical clustering (96.8% overall concordance in assignment of haplotypes to groups).

**Figure 2.**
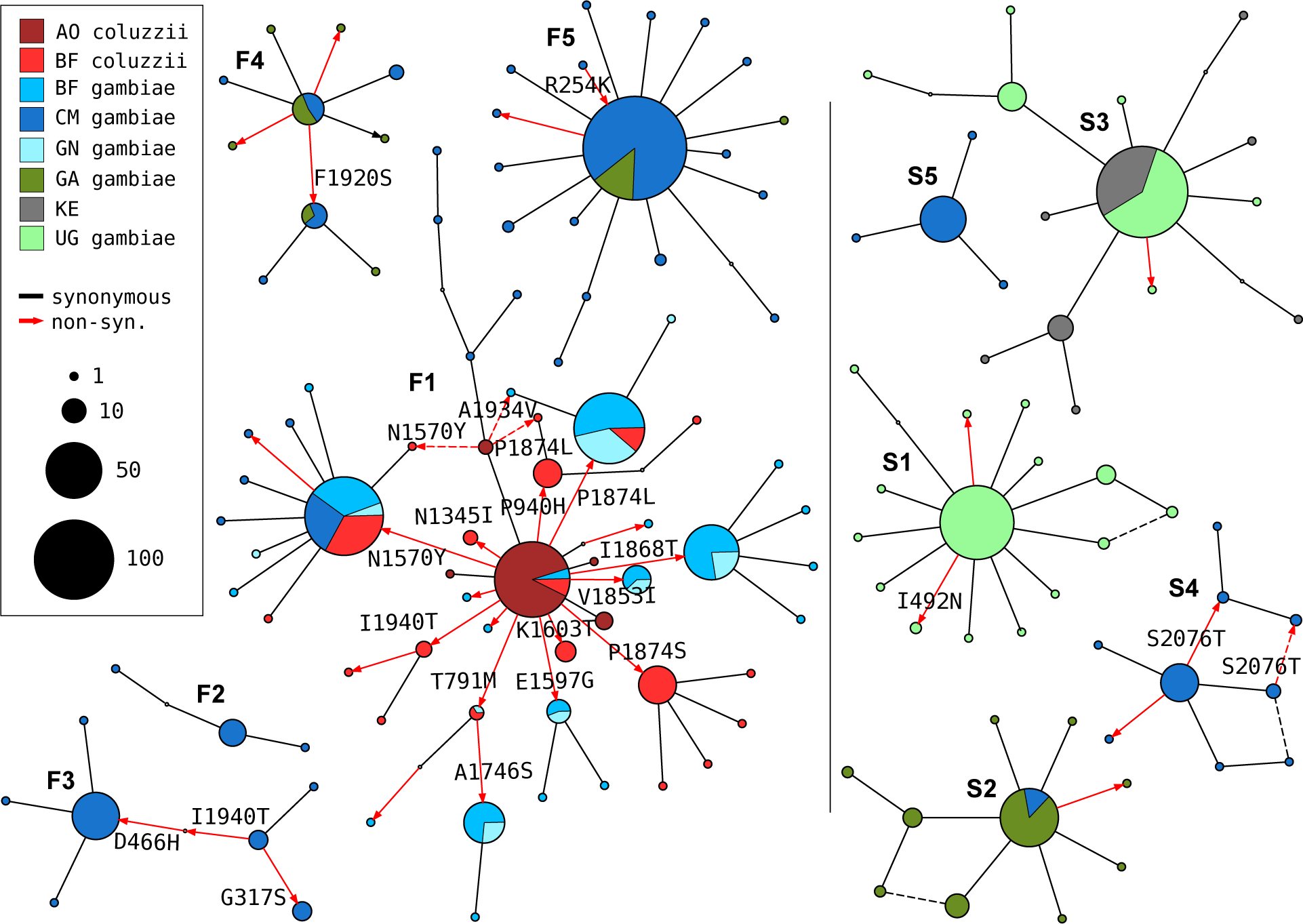
Haplotype networks. Median joining network for haplotypes carrying `L995F` (labelled F1-F5) or `L995S` variants (S1-S5) with a maximum edge distance of two SNPs. Labelling of network components is via concordance with hierarchical clusters discovered in [19]. Node size is relative to the number of haplotypes contained and node colour represents the proportion of haplotypes from mosquito populations/species -AO=Angola; BF=Burkina Faso; GN=Guinea; CM=Cameroon; GA=Gabon; UG=Uganda; KE=Kenya. Non-synonymous edges are highlighted in red and those leading to non-singleton nodes are labelled with the codon change, arrow head indicates direction of change away from the reference allele. Network components with fewer than three haplotypes are not shown.

The haplotype network brings into sharp relief the explosive radiation of amino acid substitutions secondary to the `L995F` allele (Figure 2). Within the F1 group, nodes carrying non-synonymous variants radiate out from a central node carrying only `L995F`, suggesting that the central node represents the ancestral haplotype carrying `L995F` alone which initially came under selection, and these secondary variants have arisen subsequently as new mutations. Many of the nodes carrying secondary variants are large, consistent with positive selection and a functional role for these secondary variants as modifiers of the `L995F` resistance phenotype. The F1 network also allows us to infer multiple introgression events between the two species. The central (putatively ancestral) node contains haplotypes from individuals of both species, as do nodes carrying the `N1570Y`, `P1874L` and `T791M` variants. This structure is consistent with an initial introgression of the ancestral F1 haplotype, followed later by introgressions of haplotypes carrying secondary mutations. The haplotype network also illustrates the constrasting levels of non-synonymous variation between `L995F` and `L995S`. Only two non-synonymous variants are present within the `L995S` groups, and both are at low frequency, thus may be neutral or mildly deleterious variants that are hitch-hiking on selective sweeps for the `L995S` allele.

The F1 group contained haplotypes from mosquitoes of both species, and from mosquitoes sampled in 4 different countries (Guinea, Burkina Faso, Cameroon, Angola) (Figure 3). The F4, F5 and S2 groups each contained haplotypes from both Cameroon and Gabon. The S3 group contained haplotypes from both Uganda and Kenya. The haplotypes within each of these groups were nearly identical across the entire span of the *Vgsc* gene (*π <* 5.1 *×* 10^*−*5^ *bp^−^*^1^). In contrast, diversity among wild-type haplotypes was two orders of magnitude greater (Cameroon *An. gambiae π* = 1.4 *×* 10^*−*3^ *bp^−^*^1^; Guinea-Bissau *π* = 5.7 *×* 10^*−*3^ *bp^−^*^1^). Thus it is reasonable to assume that each of these five groups contains descendants of an ancestral haplotype that carried a resistance allele and has risen in frequency due to selection for insecticide resistance. Given this assumption, these groups each provide evidence for adaptive gene flow between mosquito populations separated by considerable geographical distances.

**Figure 3.**
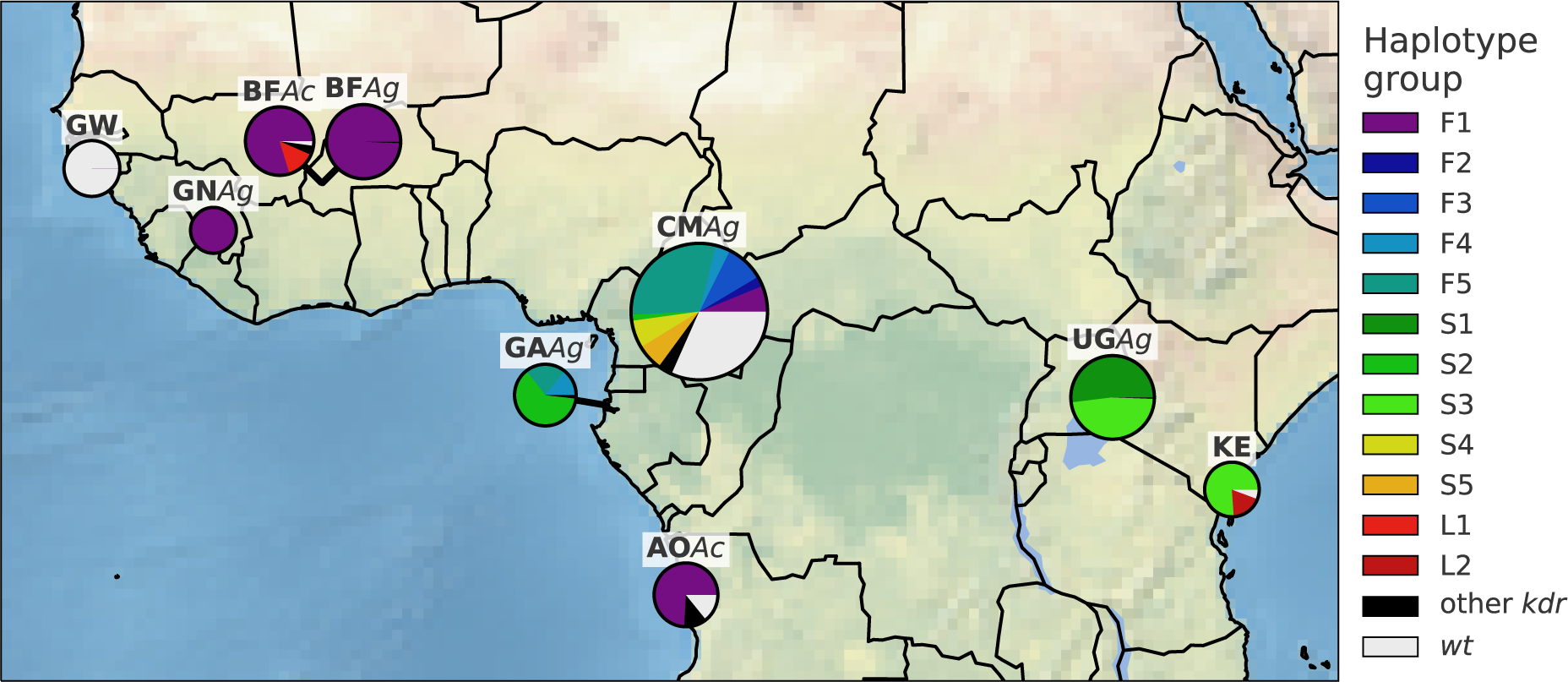
Map of haplotype frequencies. Each pie shows the frequency of different haplotype groups within one of the populations sampled. The size of the pie is proportional to the number of haplotypes sampled. The size of each wedge within the pie is proportional to the frequency of a haplotype group within the population. Haplotypes in groups F1-5 carry the `L995F` *kdr* allele. Haplotypes in groups S1-5 carry the `L995S` *kdr* allele. Haplotypes in group L1 carry the I1527T allele. Haplotypes in group L2 carry the M490I allele. Wild-type (*wt*) haplotypes do not carry any known or putative resistance alleles.

A limitation of both the hierarchical clustering and network analyses is that they rely on genetic distances within a fixed genomic window from the start to the end of the *Vgsc* gene. *Anopheles* mosquitoes undergo homologous recombination during meiosis in both males and females, and any recombination events that occurred within this genomic window could affect the way that haplotypes are grouped together in clusters or network components. In particular, recombination events could occur during the geographical spread of a resistance allele, altering the genetic background upstream and/or downstream of the allele itself. An analysis based on a fixed genomic window might then fail to infer gene flow between two mosquito populations, because haplotypes with and without a recombination event could be grouped separately, despite the fact that they share a recent common ancestor. To investigate the possibility that recombination events may have affected our grouping of haplotypes carrying resistance alleles, we performed a windowed analysis of haplotype homozygosity, spanning *Vgsc* and up to a megabase upstream and downstream of the gene (Supplementary Figures S1, S2). This analysis supported a refinement of our initial grouping of haplotypes carrying resistance alleles. All haplotypes within groups S4 and S5 were effectively identical on both the upstream and downstream flanks of the gene, but there was a region of divergence within the *Vgsc* gene itself that separated them in the fixed window analyses (Supplementary Figure S2). The 13.8 kbp region of divergence occurred upstream of codon 995 and contained 8 SNPs that were fixed differences between S4 and S5. A possible explanation for this short region of divergence is that a gene conversion event has occurred within the gene, bringing a segment from a different genetic background onto the original genetic background on which the `L995S` resistance mutation occurred.

### Positive selection for resistance alleles

To investigate evidence for positive selection on non-synonymous alleles, we performed an analysis of extended haplotype homozygosity (EHH) [25]. Haplotypes under recent positive selection will have increased rapidly in frequency, thus have had less time to be broken down by recombination, and should on average have longer regions of haplotype homozygosity relative to wild-type haplotypes. We defined a core region spanning *Vgsc* codon 995 and an additional 6 kbp of flanking sequence, which was the minimum required to differentiate the haplotype groups identified via clustering and network analyses. Within this core region, we found 18 distinct haplotypes at a frequency above 1% within the cohort. These included core haplotypes corresponding to each of the 10 haplotype groups carrying `L995F` or `L995S` alleles identified above, as well as a core haplotype carrying `I1527T` which we labelled L1 (due to it carrying the the wild-type leucine codon at position 995). We also found a core haplotype corresponding to a group of haplotypes from Kenya carrying an `M490I` allele, which we labelled as L2. All other core haplotypes we labelled as wild-type (*wt*). We then computed EHH decay for each core haplotype up to a megabase upstream and downstream of the core locus (Figure 4).

**Figure 4.**
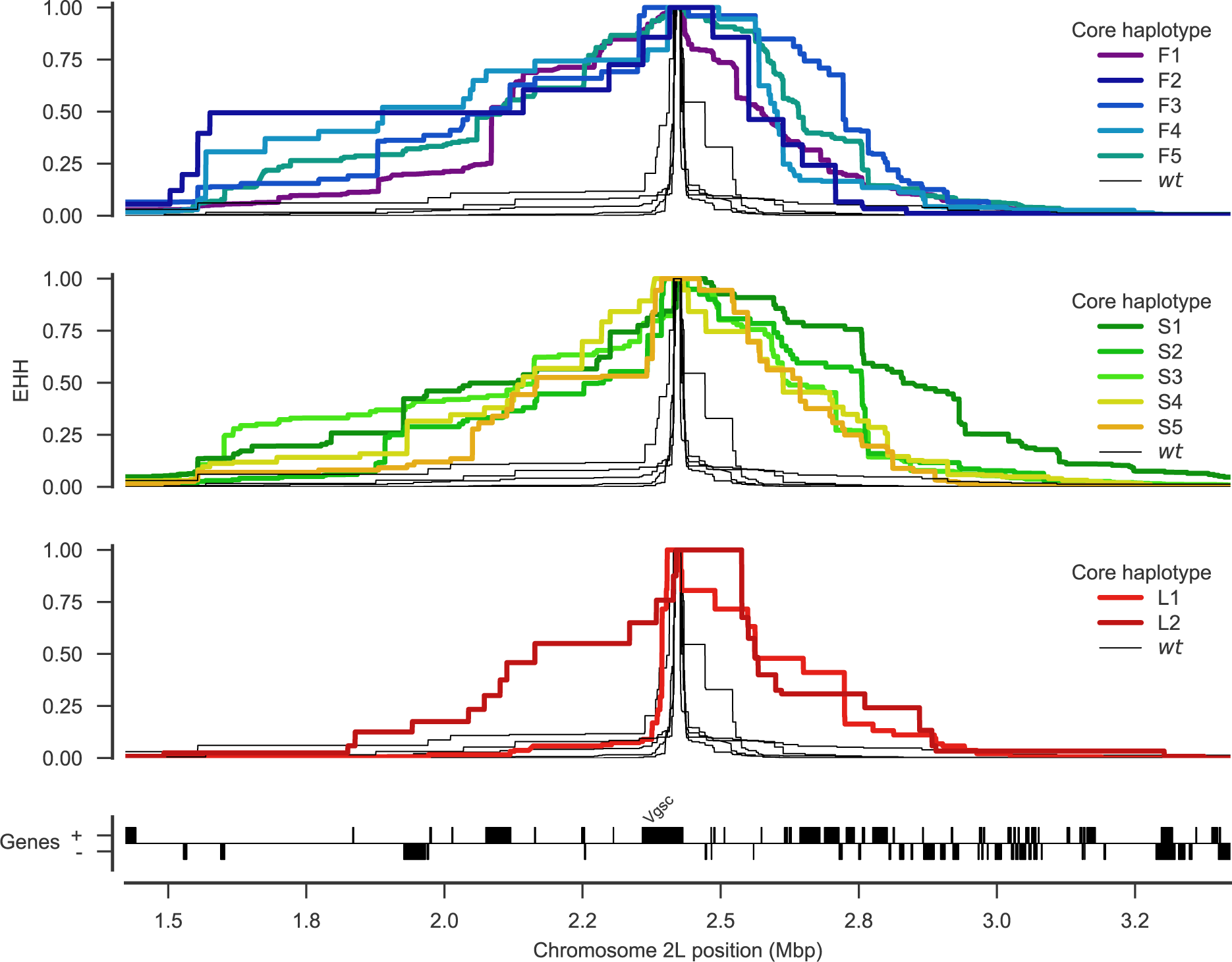
Evidence for positive selection on haplotypes carrying known or putative resistance alleles. Each panel plots the decay of extended haplotype homozygosity (EHH) for a set of core haplotypes centred on *Vgsc* codon 995. Core haplotypes F1-F5 carry the `L995F` allele; S1-S5 carry the `L995S` allele; L1 carries the I1527T allele; L2 carries the M490I allele. Wild-type (*wt*) haplotypes do not carry known or putative resistance alleles. A slower decay of EHH relative to wild-type haplotypes implies positive selection (each panel plots the same collection of wild-type haplotypes).

As expected, haplotypes carrying the `L995F` and `L995S` resistance alleles all experience a dramatically slower decay of EHH relative to wild-type haplotypes, supporting positive selection. Previous studies have found evidence for different rates of EHH decay between `L995F` and `L995S` haplotypes, suggesting differences in the timing and/or strength of selection [15]. However, we found no systematic difference in the length of shared haplotypes when comparing F1-5 (carrying `L995F`) against S1-5 (carrying L995S) (Supplementary Figure S3). There were, however, some differences between core haplotypes carrying the same allele. For example, shared haplotypes were significantly longer for S1 (median 1.091 cM, 95% CI [1.076 −1.091]) versus other core haplotypes carrying `L995S` (e.g., S2 median 0.699 cM, 95% CI [0.696 - 0.705]; Supplementary Figure S3). Longer shared haplotypes indicate a more recent common ancestor, and thus some of these core haplotypes may have experienced more recent and/or more intense selection than others. The L1 haplotype carrying `I1527T+V402L` exhibited a slow decay of EHH on the downstream flank of the gene, similar to haplotypes carrying `L995F` and `L995S`, indicating that this combination of alleles has experienced positive selection. EHH decay on the upstream gene flank was faster, being similar to wild-type haplotypes, however there were two separate nucleotide substitutions encoding `V402L` within this group of haplotypes, and a faster EHH decay on this flank is consistent with recombination events bringing `V402L` alleles from different genetic backgrounds together with an ancestral haplotype carrying `I1527T`. The L2 haplotype carrying `M490I` exhibited EHH decay on both flanks comparable to haplotypes carrying known resistance alleles. This could indicate evidence for selection on the `M490I` allele, however these haplotypes are derived from a Kenyan mosquito population where there is evidence for a severe recent bottleneck [19], and there were not enough wild-type haplotypes from Kenya with which to compare, thus this signal may also be due to the extreme demographic history of this population.

## Discussion

### Cross-resistance between pyrethroids and DDT

The VGSC protein is the physiological target of both pyrethroid insecticides and DDT [4]. The `L995F` and `L995S` alleles are known to increase resistance to both of these insecticide classes [6, 8]. By 2012, over half of African households owned at least one pyrethroid impregnated ITN and nearly two thirds of IRS programmes were using pyrethroids [2]. Pyrethroids were also introduced into agriculture in Africa prior to the scale-up of public health vector control programmes, and continue to be used on a variety of crops such as cotton [26]. DDT was used in Africa for several pilot IRS projects carried out during the first global campaign to eradicate malaria, during the 1950s and 1960s [11]. DDT is still approved for IRS use by WHO and remains in use in some locations, however within the last two decades pyrethroid use has been far more common and widespread. DDT was also used in agriculture from the 1940s, and although agricultural usage has greatly diminished since the 1970s, some usage remains [27]. In this study we reported evidence of positive selection on the `L995F` and `L995S` alleles, as well as the `I1527T+V402L` combination and possibly `M490I`. We also found 14 other non-synonymous substitutions that have arisen in association with `L995F` and appear to be positively selected. Given that pyrethroids have dominated public health insecticide use for two decades, it is reasonable to assume that the selection pressure on these alleles is primarily due to pyrethroids rather than DDT. It has previously been suggested that `L995S` may have been initially selected by DDT usage [15]. However, we did not find any systematic difference in the extent of haplotype homozygosity between these two alleles, suggesting that both alleles have been under selection over a similar time frame. We did find some significant differences in haplotype homozygosity between different genetic backgrounds carrying resistance alleles, suggesting differences in the timing and/or strength of selection these may have experienced. However, there have been differences in the scale-up of pyrethroid-based interventions in different regions, and this could in turn generate heterogeneities in selection pressures. Nevertheless, it is possible that some if not all of the alleles we have reported provide some level of cross-resistance to DDT as well as pyrethroids, and we cannot exclude the possibility that earlier DDT usage may have contributed at least in part to their selection. The differing of resistance profiles to the two types of pyrethroids (type I, e.g., permethrin; and type II, e.g., deltamethrin) [28], will also affect the selection landscape. Further sampling and analysis is required to investigate the timing of different selection events and relate these to historical patterns of insecticide use in different regions.

### Resistance phenotypes for novel non-synonymous variants

The sodium channel protein consists of four homologous domains (I-IV) each of which comprises six transmembrane segments (S1-S6) connected by intracellular and extracellular loops [5]. Two pyrethroid binding sites have been predicted within the pore-forming modules of the protein, the first (PyR1) involving residues from transmembrane segments IIS5 and IIIS6 and the internal linker between IIS4 and IIS5 (IIL45) [29], the second (PyR2) involving segments IS5, IS6, IIS6 and IL45 [22, 5]. Many of the amino acid substitutions known to cause pyrethroid resistance in insects affect residues within one of these two pyrethroid binding sites, and thus can directly alter pyrethroid binding [5]. For example, the `L995F` and `L995S` substitutions occur in segment IIS6 and belong to binding site PyR2 [22]. The `I1527T` substitution that we discovered in *An. coluzzii* mosquitoes from Burkina Faso occurs in segment IIIS6 and is immediately adjacent to two pyrethroid-sensing residues in site PyR1 [5]. It is thus plausible that pyrethroid binding could be altered by this substitution. The I1527T substitution (*M. domestica* codon 1532) has been found in *Aedes albopictus* [30], and substitutions in the nearby codon 1529 (*M. domestica* codon 1534) have been reported in *Aedes albopictus* and in *Aedes aegypti* where it was found to be associated with pyrethroid resistance [5, 31, 32]. We found the `I1527T` allele in tight linkage with two alleles causing a `V402L` substitution (*M. domestica* codon 410). Substitutions in codon 402 have been found in multiple insect species and are by themselves sufficient to confer pyrethroid resistance [5]. Codon 402 is within segment IS6, immediately adjacent to a pyrethroid sensing residue in site PyR2. The fact that we find `I1527T` and `V402L` in such tight mutual association is intriguing because (a) these two residues appear to affect different pyrethroid binding sites, and (b) haplotypes carrying `V402L` alone should also have been positively selected and thus be present in one or more populations.

A number of substitutions in segments of the protein that are not involved in either of the two pyrethroid binding sites have also been shown to confer pyrethroid resistance. For example, the `N1570Y` substitution causes substantially enhanced pyrethroid resistance when combined with `L995F`, although codon 1570 occurs in the internal linker between domains III and IV (LIII/IV) [22]. Computer modelling of the protein structure has suggested that substitutions in codon 1570 could allosterically alter site PyR2 and thus affect pyrethroid binding [22]. In addition to `N1570Y`, we found thirteen other substitutions at appreciable frequency occurring almost exclusively in association with `L995F` (Table 1; Figure 1). Of these, two (D466H, E1597G) occurred in the larger internal linkers between protein domains, one (R254K) occurred within a smaller internal linker between domain subunits, two (T791M, K1603T) occurred within an outer (“voltage-sensing”) transmembrane segment, one (A1746S) occurred within an inner (“pore-forming”) transmembrane segment, and the remaining seven occurred in the internal carboxyl-terminal tail. Thus there is no simple pattern regarding where these variants occur within the protein structure. Further work is required to confirm which of these substitutions affect pyrethroid resistance, and to determine whether they allosterically modify a pyrethroid binding site in a similar vein to `N1570Y`, or whether they provide some other benefit such as compensating for a deleterious effect of `L995F` on normal nervous system function. The novel `M490I` substitution, found on the Kenyan L2 haplotypic background potentially under selection, also occurs in an internal linker between protein domains (LI/II). However, M490I did not occur in association with `L995F` or any other non-synonymous substitutions. It is plausible that substitutions outside of pyrethroid binding sites could independently confer an insecticide resistance phenotype, because there are several known examples in other insect species [5]. Work in other species has also suggested that pyrethroid resistance substitutions could act not by altering pyrethroid binding but by altering the channel gating kinetics or the voltage-dependence of activation [5]. Thus there are a number of potential mechanisms by which a pyrethroid resistance phenotype can be obtained, and clearly much remains to be unravelled regarding the molecular biology of pyrethroid resistance in this gene.

### Design of genetic assays for surveillance of pyrethroid resistance

Entomological surveillance teams in Africa regularly genotype mosquitoes for resistance alleles in *Vgsc* codon 995, and use those results as an indicator for the presence of pyrethroid resistance alongside results from insecticide resistance bioassays. They typically do not, however, sequence the gene or genotype any other polymorphisms within the gene. Thus if there are other polymorphisms within the gene that cause or significantly enhance pyrethroid resistance, these will not be detected. Also, if a codon 995 resistance allele is observed, there is no way to know whether the allele is on a genetic background that has also been observed in other mosquito populations, and thus no way to investigate whether resistance alleles are emerging locally or being imported from elsewhere. Whole-genome sequencing of individual mosquitoes clearly provides data of sufficient resolution to answer these questions, and could be used to provide ongoing resistance surveillance. The cost of whole-genome sequencing continues to fall, with the present cost being approximately 50 GBP to obtain ~30*×* coverage of an individual *Anopheles* mosquito genome with 150 bp paired-end reads. However, to achieve substantial spatial and temporal coverage of mosquito populations, it is currently cheaper and more practical to develop targeted genetic assays for resistance outbreak surveillance. Technologies such as amplicon sequencing [33] could scale to tens of thousands of mosquitoes at low cost and could be implemented using existing platforms in national molecular biology facilities.

To facilitate the development of targeted genetic assays for surveillance of *Vgsc*-mediated pyrethroid resistance, we have produced several supplementary data tables. In Supplementary Table 1 we list all 64 non-synonymous variants found within the *Vgsc* gene in this study, with population allele frequencies. In Supplementary Table 2 we list 771 biallelic SNPs, within the *Vgsc* gene and up to 10 kbp upstream or downstream, that are potentially informative regarding which haplotype group a resistance haplotype belongs to, and thus could be used for tracking the spread of resistance. This table includes the allele frequency within each of the 12 haplotype groups defined here, to aid in identifying SNPs that are highly differentiated between two or more haplotype groups. We also provide Supplementary Table 3 which lists all 8,297 SNPs found within the *Vgsc* gene and up to 10 kbp upstream or downstream, which might need to be taken into account as flanking variation when searching for PCR primers to amplify a SNP of interest. To provide some indication for how many SNPs would need to be assayed in order to track the spread of resistance, we used haplotype data from this study to construct decision trees that could classify which of the 12 groups a given haplotype belongs to (Figure 5). This analysis suggested that it should be possible to construct a decision tree able to classify haplotypes with >95% accuracy by using 20 SNPs or less. In practice, more SNPs would be needed, to provide some redundancy, and also to type non-synonymous polymorphisms in addition to identifying the genetic background. However, it is still likely to be well within the number of SNPs that could be assayed in a single multiplex via amplicon sequencing. Thus it should be feasible to produce low-cost, high-throughput genetic assays for tracking the spread of pyrethroid resistance. If combined with a limited amount of whole-genome sequencing at sentinel sites, this should also allow the identification of newly emerging resistance outbreaks.

**Figure 5.**
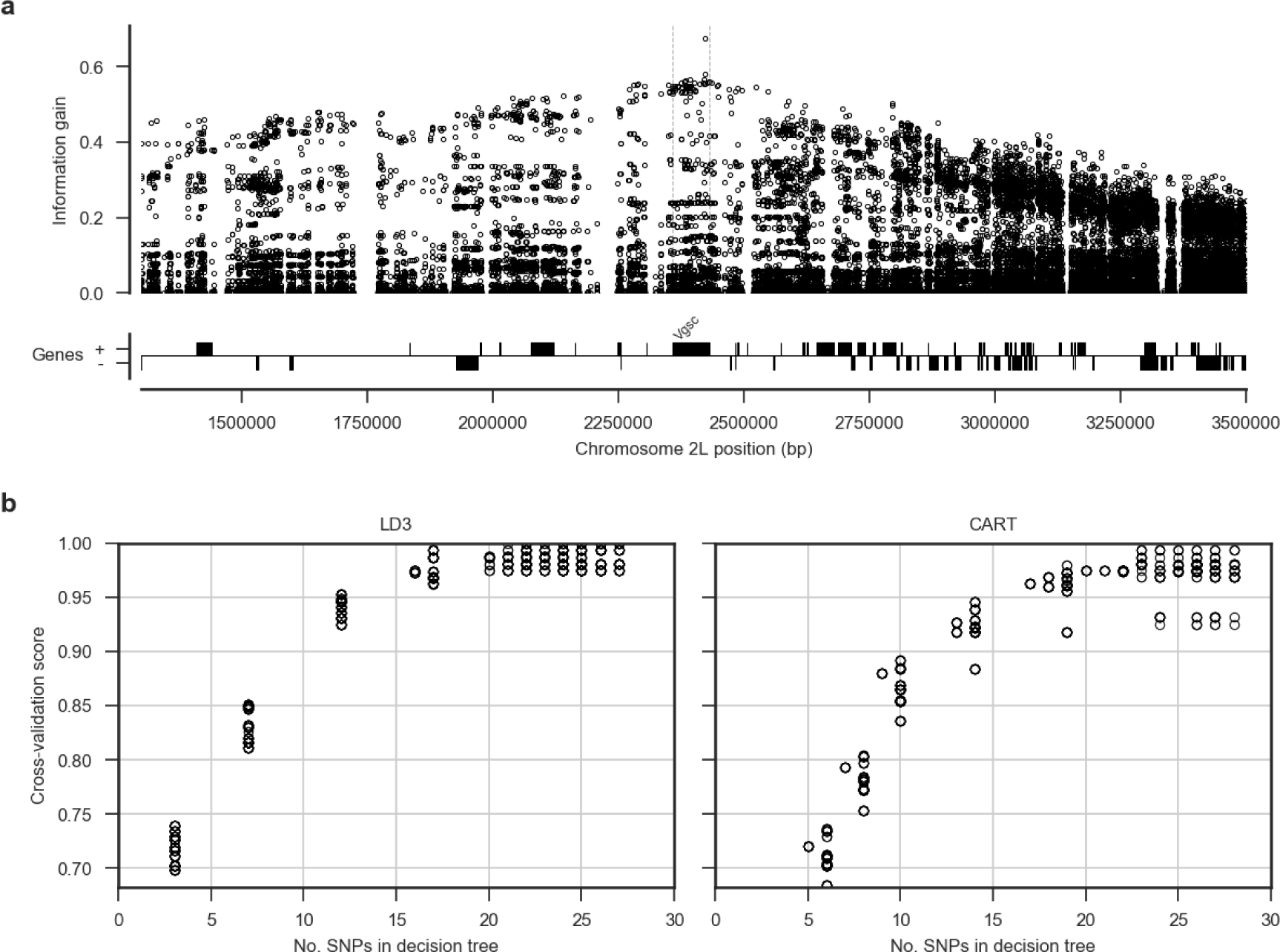
Informative SNPs for haplotype surveillance. **a**, Each data point represents a single SNP. The information gain value for each SNP provides an indication of how informative the SNP is likely to be if used as part of a genetic assay for testing whether a mosquito carries a resistance haplotype, and if so, which haplotype group it belongs to. **b**, Number of SNPs required to accurately predict which group a resistance haplotype belongs to. Each data point represents a single decision tree. Decision trees were constructed using either the LD3 (left) or CART (right) algorithm for comparison. Accuracy was evaluated using 10-fold stratified cross-validation.

## Methods

### Code

All scripts and Jupyter Notebooks used to generate analyses, figures and tables are available from the GitHub repository https://github.com/malariagen/agam-vgsc-report.

### Data

We used variant calls from the Ag1000G Phase 1 AR3 data release (https://www.malariagen.net/data/ag1000g-phase1-ar3) and phased haplotype data from the Ag1000G Phase 1 AR3.1 data release (https://www.malariagen.net/data/ag1000g-phase1-ar3.1). Variant calls from Ag1000G Phase 1 are also available from the European Nucleotide Archive (ENA; http://www.ebi.ac.uk/ena) under study PRJEB18691.

### Data collection and processing

For detailed information on Ag1000G WGS sample collection, sequencing, variant calling, quality control and phasing, see [19]. In brief, *An. gambiae* and *An. coluzzii* mosquitoes were collected from eight countries across Sub-Saharan Africa: Angola, Burkina Faso, Cameroon, Gabon, Guinea, Guinea Bissau, Kenya and Uganda. From Angola just *An. coluzzii* were sampled, Burkina Faso had samples of both *An. gambiae* and *An. coluzzii* and all other populations consisted of purely *An. gambiae*, except for Kenya and Guinea Bissau where species status is uncertain [19]. Mosquitoes were individually whole genome sequenced on the Illumina HiSeq 2000 platform, generating 100bp paired-end reads. Sequence reads were aligned to the *An. gambiae* AgamP3 reference genome assembly [34]. Aligned bam files underwent improvement, before variants were called using GATK UnifiedGenotyper. Quality control included removal of samples with mean coverage <= 14x and filtering of variants with attributes that were correlated with Mendelian error in genetic crosses.

The Ag1000G variant data was functionally annotated using the SnpEff v4.1b software [35]. Non-synonymous *Vgsc* variants were identified as all variants in transcript AGAP004707-RA with a SnpEff annotation of “missense”. The *Vgsc* gene is known to exhibit alternative splicing [4], however at the time of writing the *An. gambiae* gene annotations did not include the alternative transcripts reported by Davies et al. We wrote a Python script to check for the presence of variants that are synonymous according to transcript AGAP004707-RA but non-synonymous according to one of the other transcripts present in the gene annotations or in the set reported by Davies et al. Supplementary Table 1 includes the predicted effect for all SNPs that are non-synonymous in one or more of these transcripts. None of the variants that are non-synonymous in a transcript other than AGAP004707-RA were found to be above 5% frequency in any population.

For ease of comparison with previous work on *Vgsc*, pan Insecta, in Table 1 and Supplementary Table 1 we report codon numbering for both *An. gambiae* and *Musca domestica* (the species in which the gene was first discovered). The *M. domestica Vgsc* sequence (EMBL accession X96668 [9]) was aligned with the *An. gambiae* AGAP004707-RA sequence (AgamP4.4 gene-set) using the Mega v7 software package [36]. A map of equivalent codon numbers between the two species for the entire gene can be download from the MalariaGEN website (https://www.malariagen.net/sites/default/files/content/blogs/domestica_gambiae_map.txt).

Haplotypes for each chromosome of each sample were estimated (phased) using using phase informative reads (PIRs) and SHAPEIT2 v2.r837 [37], see [19] supplementary text for more details. The SHAPEIT2 algorithm is unable to phase multi-allelic positions, therefore the two multi-allelic non-synonymous SNPs within the *Vgsc* gene, altering codons V402 and M490, were phased onto the biallelic haplotype scaffold using MVNcall v1.0 [38]. Conservative filtering applied to the genome-wide callset had removed one of the three known insecticide resistance conferring kdr variants, `N1570Y` [10]. Manual inspection of the read alignment revealed that the SNP call could be confidently made, and it was added back into the data set and then also phased onto the haplotypes using MVNcall. Lewontin’s *D^/^* [39] was used to compute the linkage disequilibrium (LD) between all pairs of non-synonymous *Vgsc* mutations.

### Haplotype networks

Haplotype networks were constructed using the median-joining algorithm [24] as implemented in a Python module available from https://github.com/malariagen/agam-vgsc-report. Haplotypes carrying either `L995F` or `L995S` mutations were analysed with a maximum edge distance of two SNPs. Networks were rendered with the Graphviz library and a composite figure constructed using Inkscape. Non-synonymous edges were highlighted using the SnpEff annotations [35].

### Positive selection

Core haplotypes were defined on a 6,078 bp region spanning *Vgsc* codon 995, from chromosome arm 2L position 2,420,443 and ending at position 2,426,521. This region was chosen as it was the smallest region sufficient to differentiate between the ten genetic backgrounds carrying either of the known resistance alleles `L995F` or L995S. Extended haplotype homozygosity (EHH) was computed for all core haplotypes as described in [25] using scikit-allel version 1.1.9 [40], excluding non-synonymous and singleton SNPs. Analyses of haplotype homozygosity in moving windows (Supplementary Figs. S1, S2) and pairwise haplotype sharing (Supplementary Figure S3) were performed using custom Python code available from https://github.com/malariagen/agam-vgsc-report.

### Design of genetic assays for surveillance of pyrethroid resistance

To explore the feasibility of indentifying a small subset of SNPs that would be sufficient to identify each of the genetic backgrounds carrying known or putative resistance alleles, we started with an input data set of all SNPs within the *Vgsc* gene or in the flanking regions 20 kbp upstream and downstream of the gene. Each of the 1530 haplotypes in the Ag1000G Phase 1 cohort was labelled according to which core haplotype it carried, combining all core haplotypes not carrying known or putative resistance alleles together as a single "wild-type" group. Decision tree classifiers were then constructed using scikit-learn version 0.19.0 [41] for a range of maximum depths, repeating the tree construction process 10 times for each maximum depth with a different initial random state. The classification accuracy of each tree was evaluated using stratified 5-fold cross-validation.

## Footnotes

1 Codon numbering is given here relative to transcript AGAP004707-RA as defined in the AgamP4.4 gene annotations. A mapping of codon numbers from AGAP004707-RA to *Musca domestica*, the system in which knock-down resistance mutations were first described [9], is given in Table 1.

---

**Figure S1.**
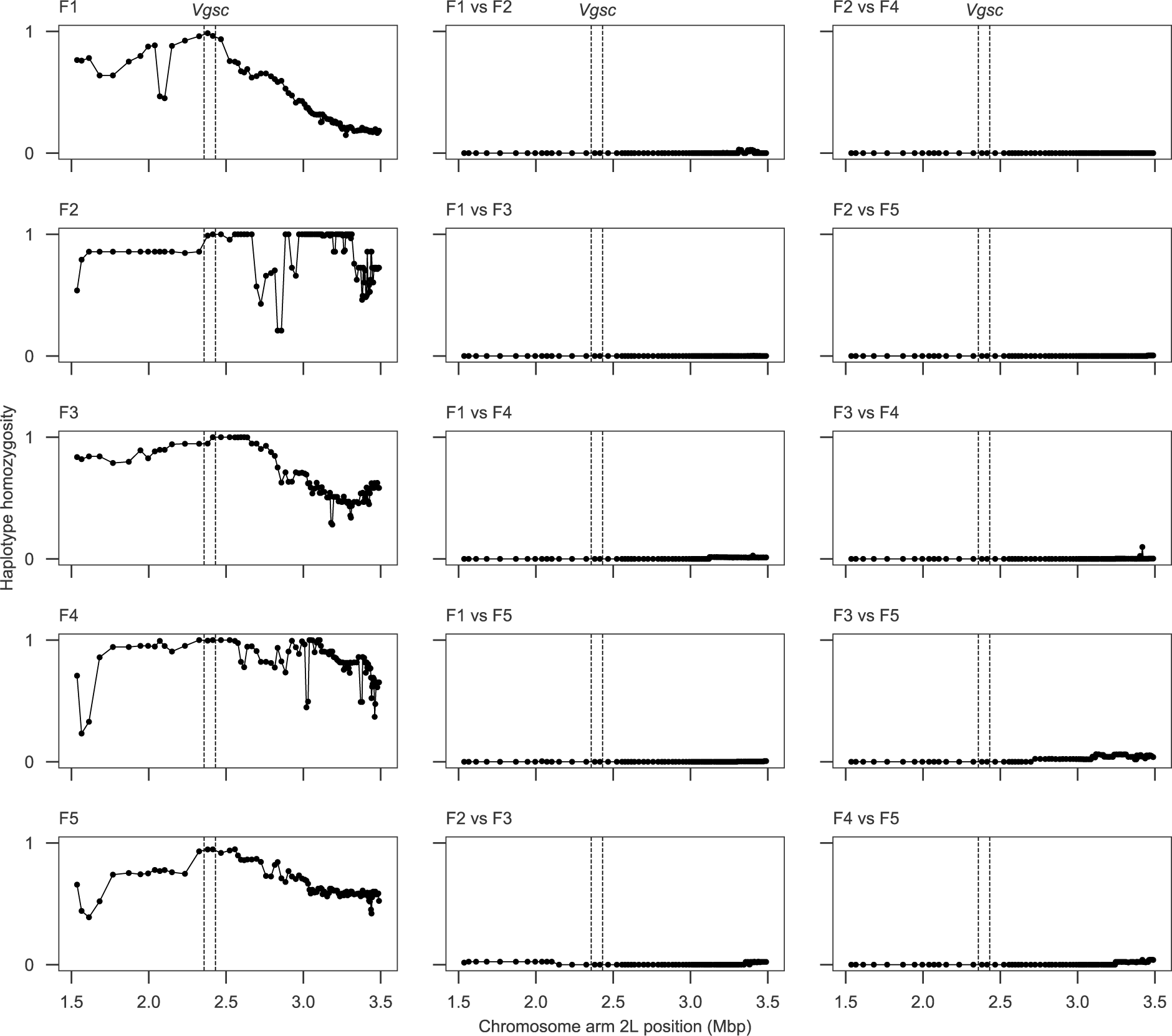
Windowed analysis of haplotype homozygosity for genetic backgrounds carrying the L995F allele. Each sub-plot shows the fraction of haplotype pairs that are identical within half-overlapping moving windows of 1000 SNPs. Each sub-plot in the left-hand column shows homozygosity for haplotype pairs within one of the haplotype groups identified by the network analysis. Sub-plots in the central and right-hand columns show homozygosity for haplotype pairs between two haplotype groups. If two haplotype groups are truly unrelated, haplotype homozygosity between them should be close to zero across the whole genome region. Dashed vertical lines show the location of the *Vgsc* gene.

**Figure S2.**
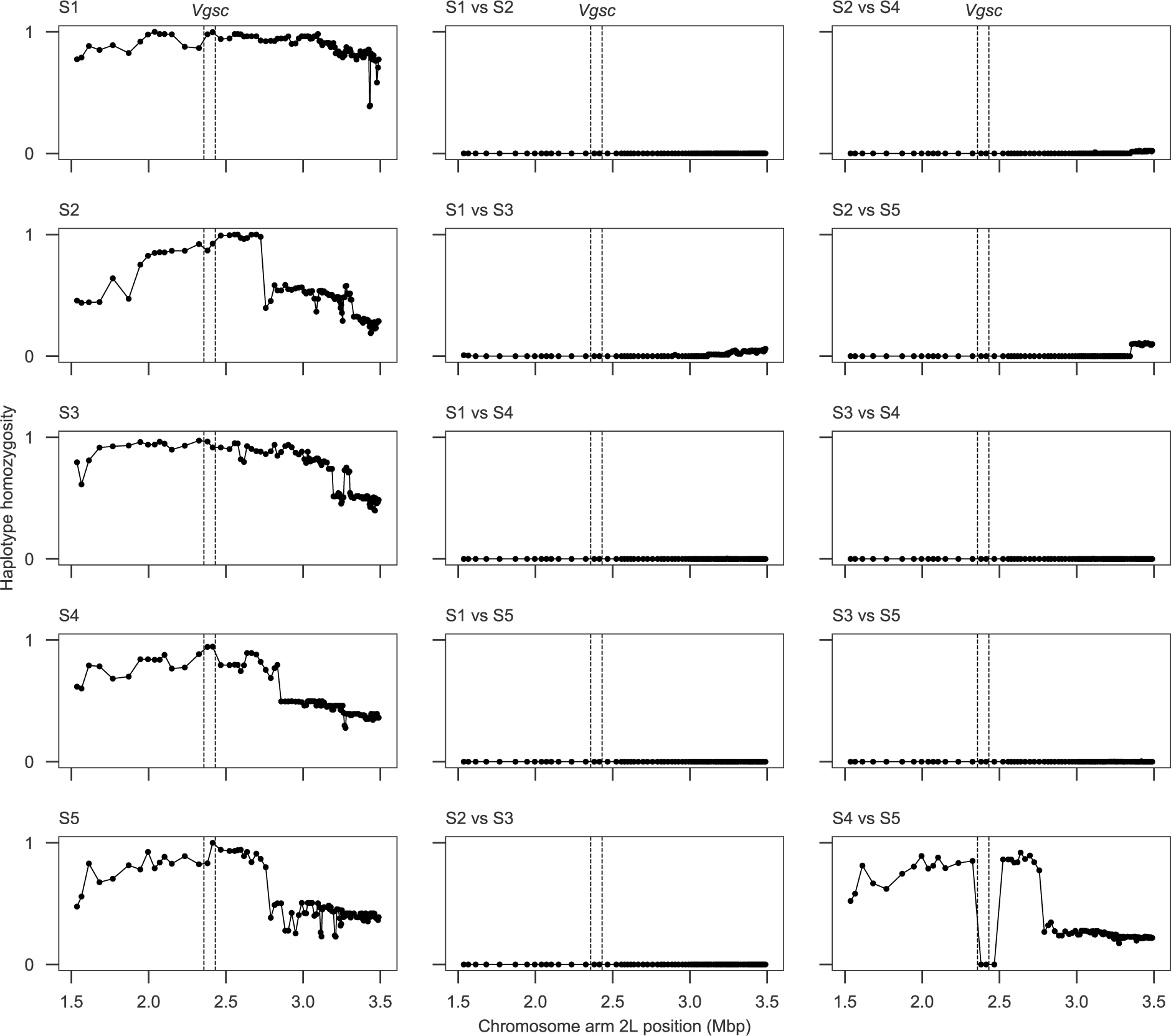
Windowed analysis of haplotype homozygosity for genetic backgrounds carrying the L995S allele. See Supplementary Figure S1 for explanation. Haplotype homozygosity is high between groups S4 and S5 on both flanks of the gene, indicating that haplotypes from both groups are in fact closely related.

**Figure S3.**
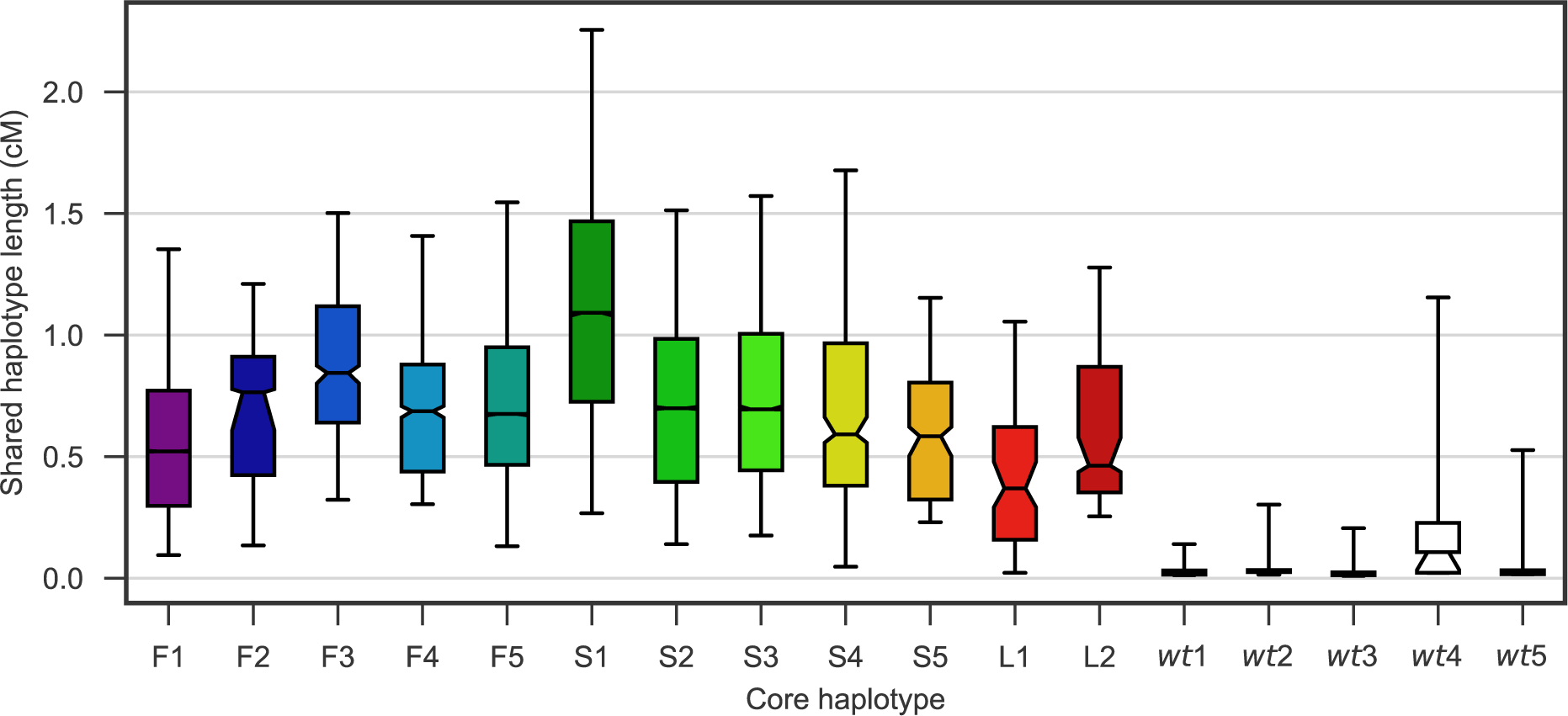
Shared haplotype length. Each bar shows the distribution of shared haplotype lengths between all pairs of haplotypes with the same core haplotype. For each pair of haplotypes, the shared haplotype length is computed as the region extending upstream and downstream from the core locus (*Vgsc* codon 995) over which haplotypes are identical at all non-singleton variants. The *Vgsc* gene sits on the border of pericentromeric heterochromatin and euchromatin, and we assume different recombination rates in upstream and downstream regions. The shared haplotype length is expressed in centiMorgans (cM) assuming a constant recombination rate of 2.0 cM/Mb on the downstream (euchromatin) flank and 0.6 cM/Mb on the upstream (heterochromatin) flank. Bars show the inter-quartile range, fliers show the 5-95th percentiles, horizontal black line shows the median, notch in bar shows the 95% bootstrap confidence interval for the median. Haplotypes F1-5 each carry the `L995F` resistance allele. Haplotypes S1-5 each carry the `L995S` resistance allele. Haplotype L1 carries the I1527T allele. Haplotype L2 carries the M490I allele. Wild-type (*wt*) haplotypes do not carry any known or putative resistance alleles.

